# Using high-resolution annotation of insect mitochondrial DNA to decipher tandem repeats in the control region

**DOI:** 10.1101/500330

**Authors:** Haishuo Ji, Xiaofeng Xu, Xiufeng Jin, Zhi Cheng, Hong Yin, Guangyuan Liu, Qiang Zhao, Ze Chen, Wenjun Bu, Shan Gao

## Abstract

In this study, we used a small RNA sequencing (sRNA-seq) based method to annotate the mitochondrial genome of the insect *Erthesina fullo* Thunberg at 1 bp resolution. Most of the new annotations were consistent with the previous annotations which were obtained using PacBio full-length transcripts. Two important findings are that animals transcribe both entire strands of mitochondrial genomes and the tandem repeat in the control region of the *E. fullo* mitochondrial genome contains the repeated Transcription Initiation Sites (TISs) of the H-strand. In addition, we found that the copy numbers of tandem repeats showed a great diversity within an individual, enriching the fundamental knowledge of mitochondrial biology. This sRNA-seq based method uses 5’ and 3’ end small RNAs to annotate nuclear non-coding and mitochondrial genes at 1 bp resolution and can also be used to identify new steady-state RNAs, particularly long non-coding RNAs (lncRNAs). Animal mitochondrial genomes containing one control region only encode two steady-state lncRNAs, which are the Mitochondrial D-loop 1 (MDL1) and its antisense gene (MDL1AS), while all other reported mitochondrial lncRNAs could be degraded fragments of transient RNAs or random breaks during experimental processing. The high-resolution annotations of mitochondrial genomes can be used to study the phylogenetics and molecular evolution of animals or to investigate mitochondrial gene transcription, RNA processing, RNA maturation and several other related topics.

## Introduction

Animal mitochondrial DNA is a small, circular, and extrachromosomal genome, typically approximately 16 kbp in size. Animal mitochondrial genomes contain 37 genes: two for rRNAs, 13 for mRNAs, 22 for tRNAs and at least one control region (CR), which is also called the Displacement-loop (D-loop) region, although the control and the D-loop region is not the same for some species [1]. The control region had not been considered as a transcriptional region until 2017, when Gao *et al*. reported that this region encodes two long non-coding RNAs (lncRNAs) [2]. The annotation of animal mitochondrial genomes usually uses blastx or structure-based covariance models [3]. Although the annotation of mitochondrial mRNAs, tRNAs and rRNAs can be easily performed using web servers (*e.g*. MITOS [3]), the annotation resolution is still limited because of the diversity in animal mitochondrial genomes, particularly the control regions. Gao *et al*. constructed the first quantitative transcription map of animal mitochondrial genomes [4] by sequencing the full-length transcriptome of the insect *Erthesina fullo* Thunberg [5] on the PacBio platform and established a straightforward and concise methodology to improve genome annotation, resulting in several findings. These findings included the 3’ polyadenylation and possible 5’ m^7^G caps of rRNAs, the polycistronic transcripts, natural antisense transcripts of mitochondrial genes [5] and two novel lncRNAs in human mitochondrial DNA [2].

Recently, 5’ and 3’ end small RNAs (5’ and 3’ sRNAs) were reported to exist in all nuclear non-coding and mitochondrial genes [6]. 5 ‘ and 3 ‘ sRNAs can be used to annotate nuclear non-coding and mitochondrial genes at 1 bp resolution and identify new steady-state RNAs, which are usually transcribed from functional genes. The high-resolution annotation of animal mitochondrial genomes can be used to investigate RNA processing, maturation, degradation, and even gene expression regulation [6]. In this study, we used this sRNA-seq based method to annotate the *E. fullo* mitochondrial genome. New findings were obtained which prove the value of high-resolution annotations of animal mitochondrial genomes and enrich the fundamental knowledge of mitochondrial biology.

## Results

### **Usi**ng 5’ and 3’ sRNAs to annotate the E. fullo mitochondrial genome

The two strands of one mitochondrial genome are differentiated by their nucleotide content. They are a guanine-rich strand referred to as the heavy strand (H-strand) and a cytosine-rich strand referred to as the light strand (L-strand). Using the sRNA-seq based method (**Materials and Methods**), we improved the annotation of the *E. fullo* mitochondrial genome (**Figure 1**) from our previous study [5]. The first important finding was that *E. fullo* transcribes both entire strands of its mitochondrial genome to produce two primary transcripts, which updated our finding that the *E. fullo* mitochondrial genome contained four transcriptional regions [5]. The quantitative transcription map of the *E. fullo* mitochondrial genome exhibited four transcriptional regions (**Figure 1A**) due to the rapid degradation of transient RNAs between these four transcriptional regions. To validate this finding, we used high-depth RNA-seq data from a strand-specific library (**Materials and Methods**) to cover 280× (4,604,253/16,485) of the *E. fullo* mitochondrial genome. As a result, all genes on both strands of the *E. fullo* mitochondrial genome were annotated (**Table 1**) to update the previous version of annotations which were obtained using PacBio full-length transcripts [4].

**Table 1.**
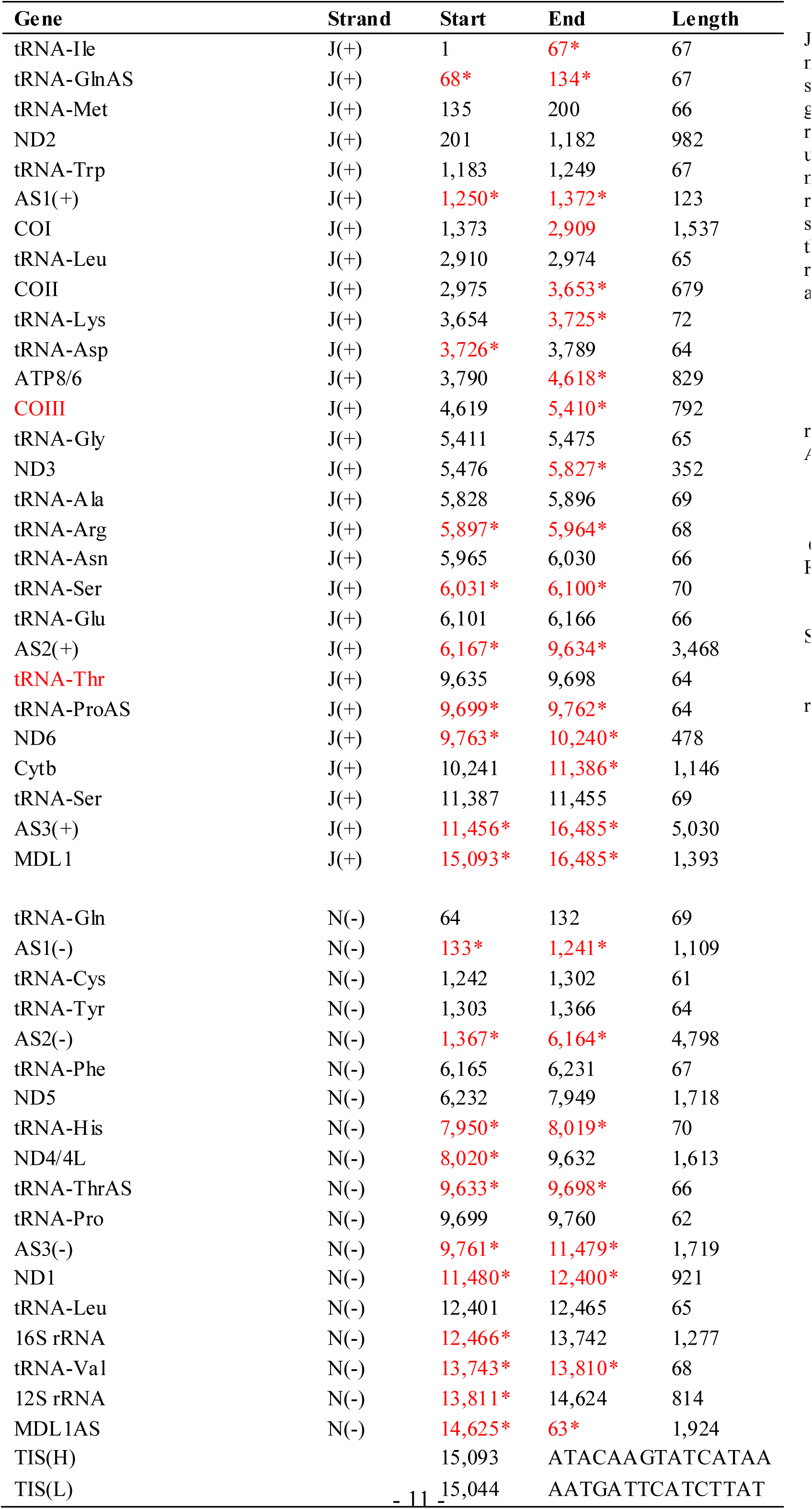
Annotation of the *E. fullo* mitochondrial genome at 1 bp resolution J(+) and N(−) represents the major and minor coding strand of the mitochondrial genome, respectively. * represents the new annotation using the sRNA-seq based method. TIS(H) and TIS(L) represents the TIS of the H-strand (TIS_H_) and the TIS of the L-strand (TIS_L_), respectively. AS1(+), AS2(+), and AS3(+) represents tRNA^Cys^AS/tRNA^Tyr^AS, tRNA^Phe^AS/ND5AS/tRNA^His^ AS/ND4/4LAS, ND1AS/tRNA^Leu^AS/16S rRNAAS/tRN A^Val^ AS/12S rRNAAS/CR, respectively. AS1(−), AS2(−), and AS3(−) represents tRNA^Met^AS/ND2AS/tRNA^Trp^ AS, COIAS/tRNA^Leu^AS/COIIAS/t RNA^Lys^AS/tRNA^Asp^AS/ATP8/6AS/COIIIAS/tRNA^Gly^AS/N D3AS/tRNA^Ala^AS/tRNA^Arg^A S/tRNA^Asn^AS/tRNA^Ser^AS/tRN A^Glu^AS, ND6AS/CytbAS/tRNA^Ser^AS, respectively.

The high-resolution annotation of the *E. fullo* mitochondrial genome was confirmed by the ‘mitochondrial cleavage’ model that we proposed in our previous study [2]. This model is based on the fact that RNA cleavage is processed: **1)** at 5’ and 3’ ends of tRNAs, **2)** between mRNAs and mRNAs except fusion genes (e.g. ATP8 and ATP6), **3)** between antisense tRNAs and mRNAs and **4)** between mRNAs and antisense tRNAs. Additionally, RNA cleavage is not processed: **1)** between mRNAs and antisense mRNAs or **2)** between antisense RNAs. In particular, this model helps the annotation of long antisense genes (*e.g*. ND1AS/tRNA^Leu^AS/16S rRNAAS/tRNA^Val^AS/12S rRNAAS/CR in *E. fullo)* in animal mitochondrial genomes, as all of them are transcribed as transient RNAs and are usually not well covered by aligned reads from sRNA-seq or RNA-seq data due to their rapid degradation.

**Figure 1.**
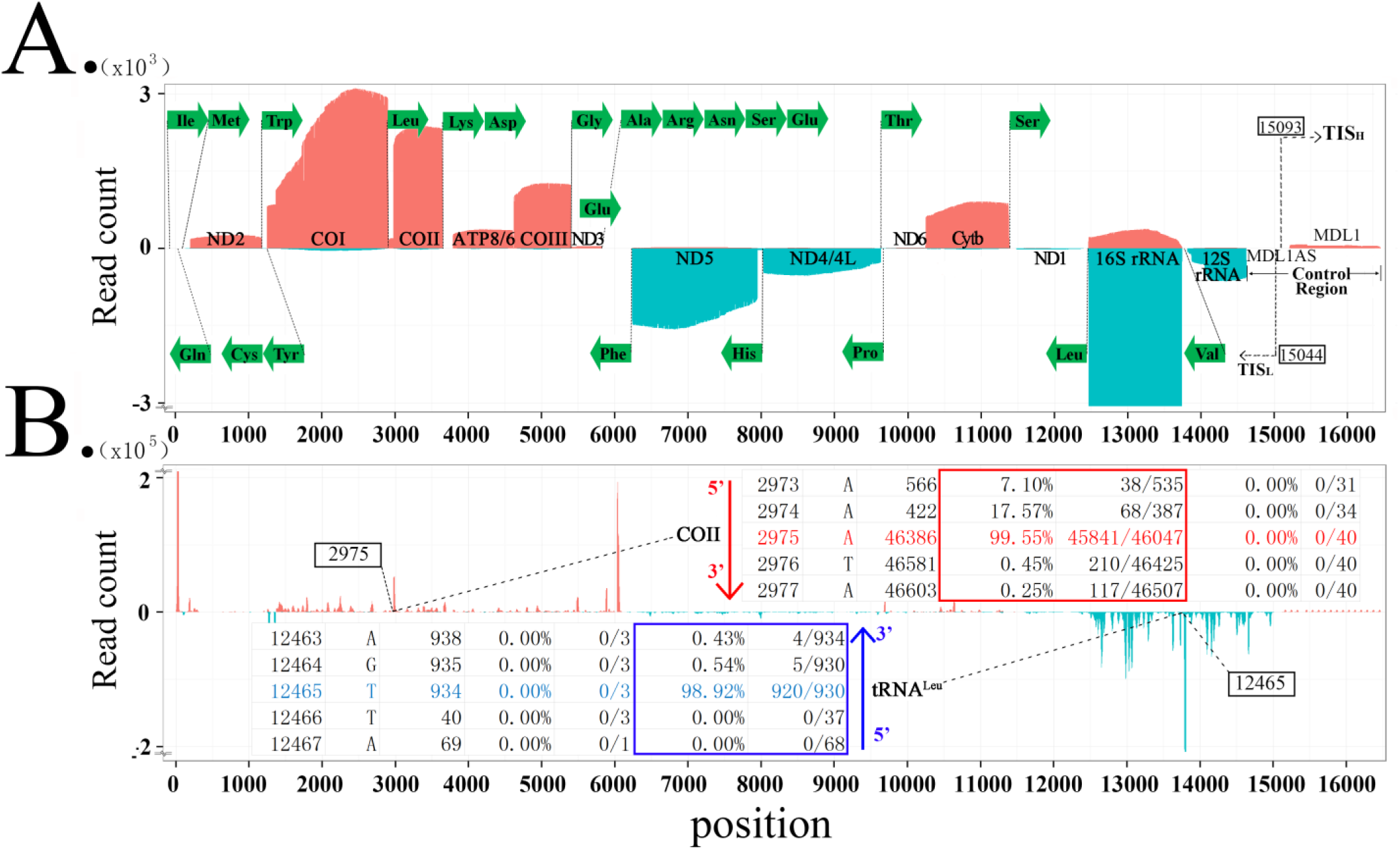
Annotation of the *E. fullo* mitochondrial genome at 1 bp resolution. **A**. Alignments of transcripts on the H-strand in red color are piled along the positive y-axis. Alignments of transcripts on the L-strand strand in blue color are piled along the negative y-axis. The tRNAs in green color are represented using their amino acids. The TIS of the H-strand (TIS_H_) and the TIS of the L-strand (TIS_L_) are at the positions 15,093 and 15,044 bp. **B**. In our previous study [6], we have defined a new file format, named “5-end format”, to easily identify the 5’ ends of mature RNAs. The 5’ enda of COII and tRNA^Leu^ are annotated at position 2,975 and 12,465 bp, respectively.

### Annotation of mitochondrial tRNAs at 1 bp resolution

The annotation resolution of mitochondrial tRNAs is limited due to the complexity of tRNA processing. Using 5’ and 3’ sRNAs, we annotated the mitochondrial tRNAs of *E. fullo* at 1 bp resolution, which can not be achieved using other existing methods. Based on these results, we propose an accurate mitochondrial tRNA processing model. One mitochondrial tRNA is cleaved from a mitochondrial primary transcript into a precursor, and then the acceptor stem of the precursor is trimmed to contain a 1 -bp overhang at the 3’ end. Finally, CCAs are post-transcriptionally added to the 3’ ends of tRNAs, one nucleotide at a time. Using other existing methods, mitochondrial tRNAs are annotated between two trimming sites of their mature RNAs, which misses the information on the cleavage sites. Using our model, mitochondrial tRNAs are annotated between two cleavage sites and the information on the trimming sites can be derived. Here, three typical examples (tRNA^Ile^, tRNA^Arg^ and tRNA^Ser^) are used to demonstrate the annotation resolution using the sRNA-seq based method (**Figure 2**).

**Figure 2.**
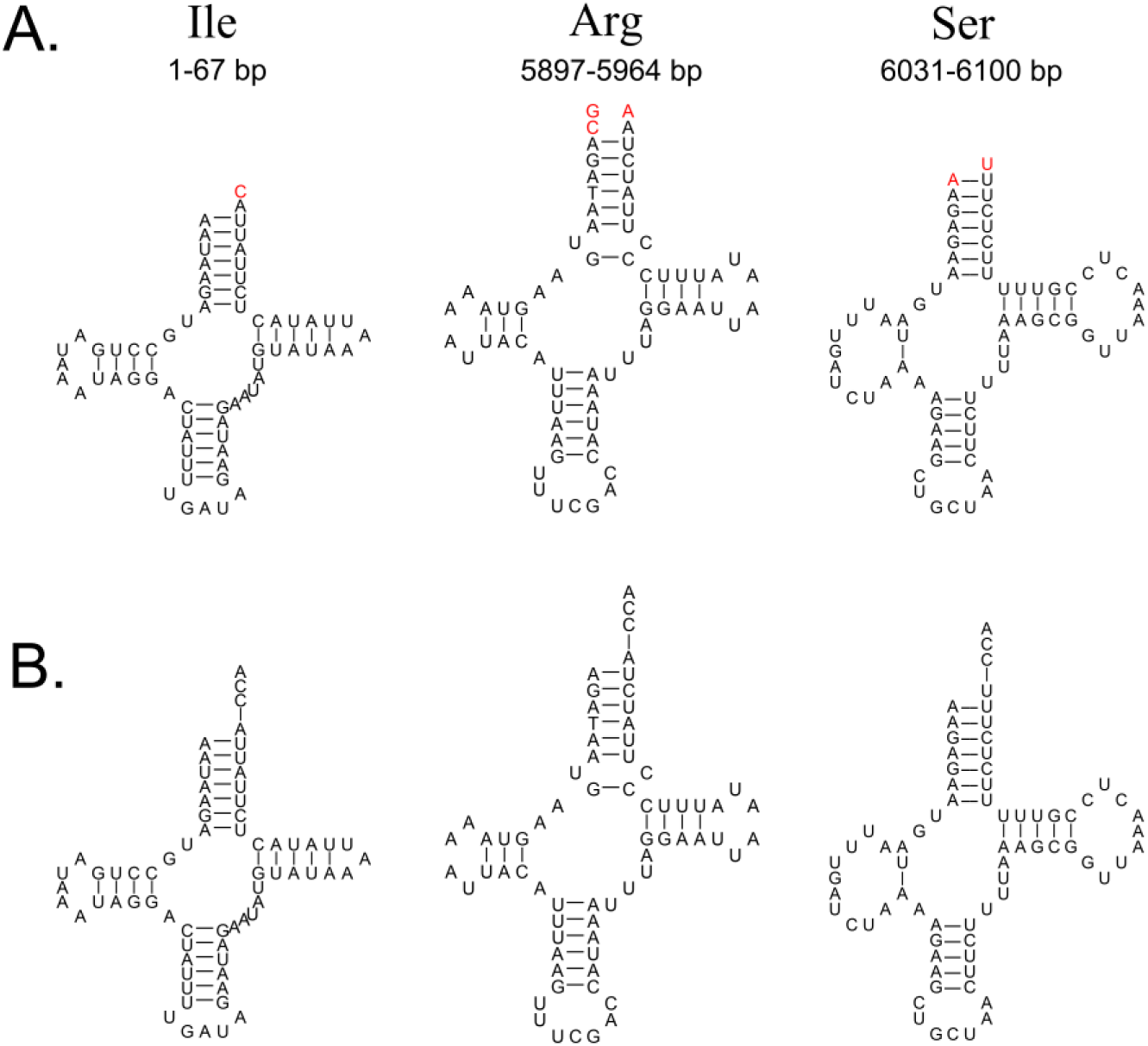
Corrected annotations of mitochondrial tRNAs. **A**. Using the sRNA-seq based method, mitochondrial tRNAs are annotated between two cleavage sites and several nucleic acids in red color are missed using other existing methods. **B**. The acceptor stem of a tRNA precursor is trimmed to contain a 1-bp overhang at the 3’ end. CCAs are post-transcriptionally added to the 3’ ends of tRNAs, one nucleotide at a time.

### The MDL1 and MDL1AS genes

Based on the human and *E. fullo* genomes, we propose that any animal mitochondrial genome that contains one control region transcribes both entire strands. However, the situation in animal mitochondrial genomes that contain more than one control region [7] is unknown. One control region contains at least two Transcription Initiation Sites (TISs), which are the TIS of the H-strand (TIS_H_) and the TIS of the L-strand (TIS_L_). The investigation of more than two TISs for one strand (*e.g*. TIS_H1_ and TIS_H2_ in human) is beyond the scope of this study. One control region is involved in at least four genomic regions (**Figure 3A**). The first and second regions that cover the complete control region are between the two nearest cleavage sites on the H-strand and L-strand, respectively. In *E. fullo*, they are ND1AS/tRNA^Leu^AS/16S rRNAAS/tRNA^Val^AS/12S rRNAAS/CR (mtDNA:11456-16485) on the H-strand and CR/tRNA^Ile^AS (mtDNA:14625-67) on the L-strand. The third region (mtDNA: 15093-16485) starts at the TIS_H_ and ends at the downstream cleavage site on the H-strand. The fourth region (mtDNA: 14625-15044) starts at the TIS_L_ and ends at the downstream cleavage site on the L-strand. As the first region overlaps the third and the second region overlaps the fourth, two of four regions can be selected to define the Mitochondrial D-loop 1 (MDL1) and its antisense gene (MDL1AS) [2]. Here, we propose the criteria for defining MDL1 and MDL1AS. If the first region from the H-strand does not span more than one control region and two antisense tRNA genes, it is defined as MDL1. Otherwise, the third region is defined as MDL1. If the second region from the L-strand does not span more than one control region and two antisense tRNA genes, it is defined as MDL1AS. Otherwise, the fourth region is defined as MDL1AS. The human MDL1 (hsa-MDL1) contains tRNA^Pro^AS and the control region, while MDL1 in *E. fullo* (eft-MDL1) starts at the TIS_H_ and ends at the end of the control region (**Figure 3A**). In humans, has-MDL1AS starts at the TIS_L_ and ends at the end of the control region, while eft-MDL1AS contains tRNA^Ile^AS and the control region (**Figure 3A**). Using 5’ and 3’ sRNAs, all the antisense genes (**Table 1**) in the *E. fullo* mitochondrial genome were identified as transient RNAs, which are not likely to perform specific functions. Therefore, we concluded that animal mitochondrial genomes containing one control region only encode two steady-state lncRNAs (MDL1 and MDL1AS), while all other reported mitochondrial lncRNAs could be degraded fragments of transient RNAs or random breaks during experimental processing [6]. This conclusion does not rule out the possibility that sRNAs degraded from transient RNAs could have detrimental effects on the regulation of gene expression [8].

### Deciphering tandem repeats in the control region

Many, but not all, control regions contain tandem repeats, which are present across a range of (mostly mammalian) species and hypothesized to be involved in the termination of transcription because of their complex secondary structures [9]. Although tandem repeats in control regions have been reported in more than 150 species, including chicken, cat, rabbit, pig, sheep, horse, Japanese monkey and human (only gastric cancer [10]), their genetic diversity and biological functions still require extensive research. In a previous study, we obtained the tandem repeats of the *E. fullo* mitochondrial genome using the PacBio full-length transcriptome data [5] and found that eft-MDL1 (mtDNA:15093-16485) was a repeating region which was composed of multiple 83-bp repeating units (noted as R) and several insert sequences (noted as A, B and so on). Most of the repeating regions detected using *E. fullo* insects in this study follow the pattern R_x_AR_y_ (**Figure 3B**) but a few follow the pattern R_x_AR_y_BR_z_, where x, y and z represent copy numbers of R. In addition, we found that TIS_H_ was at the 5’ end of the 83-bp repeating unit. This suggests that TISs are repeated in R_x_AR_y_ (x+y=n) for n times. Further study showed three frequently occurring polymorphic sites in the 83-bp repeating unit and R_x_AR_y_ was further cleaved into two RNAs (**Figure 3B**). To validate these polymorphic sites, each of short and long DNA segments in the repeating region (**Materials and Methods**) were amplified using PCR to obtain the sequences of R_x_ and R_y_ using 18 *E. fullo* adults. Sixteen short and five long segments were successfully sequenced using Sanger sequencing to determine x and y. Using our data, we found that x ranged from 1 to 3 and y ranged from 8 to 13. The total copy numbers n (x+y) were more than 10. Our hypothesis was that a large number of TISs in an individual could prevent the deleterious mutation effects and *E. fullo* insects containing a large number of TISs have an advantage in evolution. Another important finding related to tandem repeats was that the copy numbers of tandem repeats show a great diversity within an individual. This was also validated by Sanger sequencing of 16 R_x_ and 5 R_y_.

**Figure 3.**
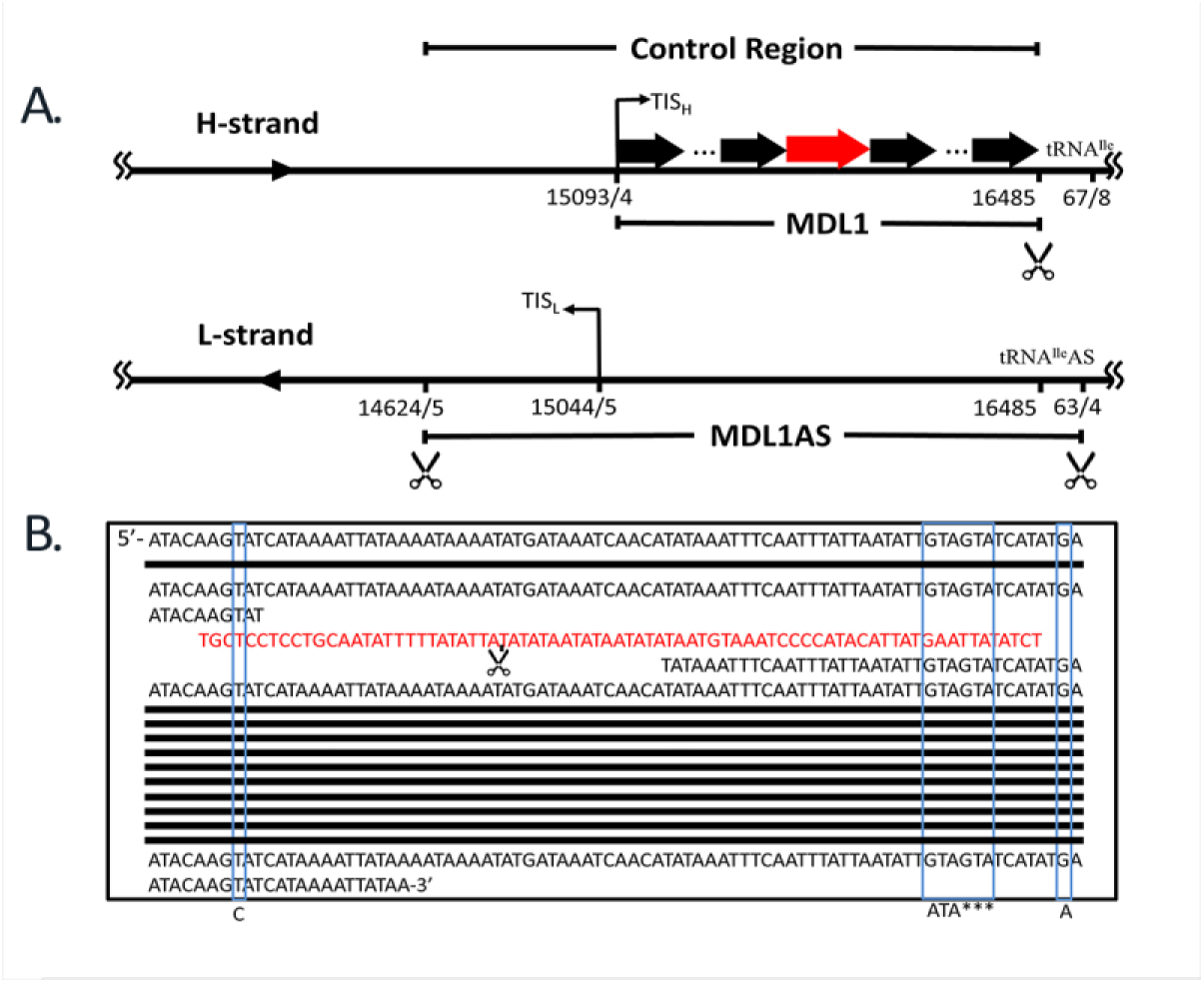
Deciphering tandem repeats in the control region. **A**. MDL1 in *E*. fullo (eft-MDL1) starts at the TIS_H_ and ends at the end of the control region, while eft-MDL1AS contains tRNA^Ile^AS and the control region. **B**. eft-MDL1 (mtDNA:15093-16485) is a repeating region which is composed of multiple 83-bp repeating units. Three frequently occurring polymorphic sites in the 83-bp repeating unit have alleles T/C, ATA/GTAGTA and G/A.

## Conclusion and discussion

In this study, we demonstrated that the sRNA-seq based method can be used to annotate mitochondrial genomes at 1 bp resolution. This improved method brought new findings, which updated our understanding of the conservation and polymorphism in mitochondrial genomes. We propose that any animal mitochondrial genome which contains one control region transcribes both entire strands. One control region contains at least one TIS_H_ and one TIS_L_, which initiate two primary transcripts. The cleavage and transcription of primary transcripts synchronize to maintain a high efficiency of mitochondrial gene expression. Animal mitochondrial genomes containing one control region only encode two steady-state lncRNAs (MDL1 and MDL1AS), while all antisense transcripts are produced from primary transcripts by cleavage but degrade rapidly.

The mechanisms for the termination of mitochondrial transcription are still unclear, as the sRNA-seq based method cannot be used to determine the Transcription Termination Sites (TTSs) of primary transcripts. Based on our findings and hypotheses, the existence of TTSs is not necessary for the transcription of mitochondrial genes and an uninterrupted transcription results in higher efficiency. In our previous study, we were interested in whether RNA polymerase had the ability to read through the TTS after the H-strand primary transcript had been completely synthesized. Surprisingly, we found two long PacBio sequences to support this "read through" model. As cells usually prefer economic and efficient ways of existence, the ‘read through’ model and the possible uninterrupted transcription of mitochondrial genomes merit further research.

As the control regions in the mitochondrial genomes are less conserved than the coding genes in evolution, they are not well investigated, compared to other mitochondrial genes. However, our findings proved that the annotations of control regions provide abundant information for the study of the phylogenetics and molecular evolution of animals. The discovery of repeated TISs suggests that control regions contain DNA elements which are highly conserved in evolution. Future work needs to be performed to investigate the copy number distribution of tandem repeats in the *E. fullo* mitochondrial genome using insects from a wide distribution. The diversity of the copy numbers within an individual can also be used to study insect development and aging.

## Materials and Methods

The PacBio full-length transcriptome data was collected from our previous study [4]. The RNA-seq and sRNA-seq data were obtained by sequencing one and three libraries, respectively. Total RNA was isolated from thoracic muscles of three, one and one *E. fullo* adults to construct three sRNA-seq libraries, respectively, which were sequenced two times as technical replicates using 50-bp single-end strategy on the Illumina HiSeq 2500 platform. Total RNA was isolated from thoracic muscles of one *E. fullo* adult to construct one RNA-seq library, which was sequenced two times as technical replicates using 150-bp paired-end strategy on the Illumina HiSeq X Ten platform. RNA-seq and sRNA-seq libraries were constructed following the protocols in our previous studies ([11] and [12]). Finally, Six runs of sRNA-seq data and four runs of RNA-seq data were obtained for this study. These data are available at the NCBI SRA database under the project accession number SRPxxxxxx.

The cleaning and quality control of sRNA-seq and RNA-seq were performed using the pipeline Fastq_clean [13] that was optimized to clean the raw reads from Illumina platforms. 291,941,849 cleaned sRNA-seq reads and 164,346,320 cleaned RNA-seq reads were used to perform and validate the new annotation of the *E. fullo* mitochondrial genome. Using the software bowtie, sRNA-seq and RNA-seq reads were aligned to the *E. fullo* mitochondrial genome with one mismatch and two mismatches, receptively. Statistics and plotting were conducted using the software R v2.15.3 the Bioconductor packages [14]. In our previous study, the annotation of *E. fullo* mitochondrial mRNAs and rRNAs was performed using the full-length transcripts and the annotation of tRNAs was performed using MITOS server [3]. The complete mitochondrial genome sequence of *E. fullo* with the new annotation using the sRNA-seq based method [6] is available at the NCBI GenBank database under the accession number xxxxxx.

The first DNA segment (mtDNA:14779-15176), named the short DNA segment, used the forward and reverse primers CTATTCCTAGCTCACATTTAAGTTCG and TGCAGGAGGAGCAATACTTG to obtain the sequence of R_x_ and the second DNA segment (mtDNA:14779-156), named the long segment, used the forward and reverse primers CTATTCCTAGCTCACATTTAAGTTCG and CCAGGGTATGAACCTGTTAGC to obtain the sequence of R_y_. The PCR reaction mixture for the first segment was incubated at 94°C for 3 min, followed by 40 PCR cycles (10 s at 94°C, 10 s at 49°C and 20 s at 72°C for each cycle) using 2×Es Taq MasterMix (CWBIO, China). The PCR reaction mixture for the second segment was incubated at 98°C for 3 min, followed by 40 PCR cycles (10 s at 98°C, 30 s at 52°C and 3 min at 72°C for each cycle) using LA Taq (TaKaRa, China).

## Acknowledgments

We appreciate the help equally from the people listed below. They are Professor Defu Chen, Bingjun He, Guoqing Liu, Dawei Huang and graduate student Siyu Li from College of Life Sciences, Nankai University.

## REFERENCES

1. J.L. Boore, Animal mitochondrial genomes. Nucleic Acids Research, 1999. 27(8): p. 1767–1780.

2. S. Gao, X. Tian, H. Chang, Y. Sun, Z. Wu, et al., Two novel lncRNAs discovered in human mitochondrial DNA using PacBio full-length transcriptome data. Mitochondrion, 2017.

3. M. Bernt, A. Donath, F. Jühling, F. Externbrink, C. Florentz, et al., MITOS: improved de novo metazoan mitochondrial genome annotation. Molecular Phylogenetics & Evolution, 2013. 69(2): p. 313–319.

4. S. Gao, Y. Ren, Y. Sun, Z. Wu, J. Ruan, et al., PacBio Full-length transcriptome profiling of insect mitochondrial gene expression. RNA biology, 2016. 13(9): p. 820–825.

5. Y. Ren, Z. Jiaqing, Y. Sun, Z. Wu, J. Ruan, et al., Full-length transcriptome sequencing on PacBio platform (in Chinese). Chinese Science Bulletin, 2016. 61(11): p. 1250–1254.

6. X. Xu, H. Ji, Z. Cheng, X. Jin, X. Yao, et al., Using pan RNA-seq analysis to reveal the ubiquitous existence of 5' end and 3' end small RNAs. bioRxiv, 2018.

7. Y. Kumazawa, H. Ota, M. Nishida and T. Ozawa, The complete nucleotide sequence of a snake (Dinodon semicarinatus) mitochondrial genome with two identical control regions. Genetics, 1998. 150(1): p. 313.

8. J. Houseley and D. Tollervey, The Many Pathways of RNA Degradation. Cell, 2009. 136(4): p. 763–776.

9. D.H. Lunt, L.E. Whipple and B.C. Hyman, Mitochondrial DNA variable number tandem repeats (VNTRs): utility and problems in molecular ecology. Molecular Ecology, 2010. 7(11): p. 1441–1455.

10. W.Y. Hung, J.C. Lin, L.M. Lee, C.W. Wu, L.M. Tseng, et al., Tandem duplication/triplication correlated with poly-cytosine stretch variation in human mitochondrial DNA D-loop region. Mutagenesis, 2008. 23(2): p. 137–142.

11. Y. Xu, S. Gao, Y. Yang, M. Huang, L. Cheng, et al., Transcriptome sequencing and whole genome expression profiling of chrysanthemum under dehydration stress. BMC Genomics, 2013. 14(1): p. 662.

12. Z. Chen, Y. Sun, X. Yang, Z. Wu, K. Guo, et al., Two featured series of rRNA-derived RNA fragments (rRFs) constitute a novel class of small RNAs. Plos One, 2017. 12(4): p. e0176458.

13. M. Zhang, F. Zhan, H. Sun, X. Gong, Z. Fei, et al. Fastq_clean: An optimized pipeline to clean the Illumina sequencing data with quality control. in Bioinformatics and Biomedicine (BIBM), 2014 IEEE International Conference on. 2014. IEEE.

14. S. Gao, J. Ou and K. Xiao, R language and Bioconductor in bioinformatics applications(Chinese Edition). 2014, Tianjin: Tianjin Science and Technology Translation Publishing Ltd.

